# The N-terminal extension of Arabidopsis ARGONAUTE 1 is essential for microRNA activities

**DOI:** 10.1101/2022.09.29.510030

**Authors:** Ye Xu, Yong Zhang, Zhenfang Li, Alyssa Soloria, Savannah Potter, Xuemei Chen

## Abstract

microRNAs (miRNAs) regulate target gene expression through their ARGONAUTE (AGO) effector protein, mainly AGO1 in *Arabidopsis thaliana*. In addition to the highly conserved N, PAZ, MID and PIWI domains with known roles in RNA silencing, AGO1 contains a long, unstructured N-terminal extension (NTE) of little-known function. Here, we show that the NTE is indispensable for the functions of Arabidopsis AGO1, as a lack of the NTE leads to seedling lethality. Within the NTE, the region containing amino acids (a.a.) 91 to 189 is essential for rescuing an *ago1* null mutant. Through global analyses of small RNAs, AGO1-associated small RNAs, and miRNA target gene expression, we show that the region containing a.a. 91-189 is required for the loading of miRNAs into AGO1. Moreover, we show that reduced nuclear partitioning of AGO1 did not affect its profiles of miRNA and ta-siRNA association. Furthermore, we show that the 1-to-90a.a. and 91-to-189a.a. regions of the NTE redundantly promote the activities of AGO1 in the biogenesis of trans-acting siRNAs. Together, we report novel roles of the NTE of Arabidopsis AGO1.

## Introduction

In eukaryotes, microRNAs (miRNAs) are ~19-to-24 nucleotide (nt) long endogenous non-coding RNAs that regulate gene expression at the post-transcriptional level through sequence complementarity with target transcripts. In plants, miRNA-mediated gene silencing is essential for a broad range of biological processes, including growth, development, and responses to abiotic and biotic stresses[1,2]. The *MIRNA* (*MIR*) genes are transcribed into long primary miRNA (pri-miRNA) transcripts by RNA POLYMERASE II in a manner similar to that of protein-coding genes[3]. The pri-miRNAs are successively processed by DICER-LIKE 1 (DCL1), an RNase III family enzyme, in a base-to-loop or loop-to-base manner to produce miRNA duplexes with 2-nt overhangs at the 3′ ends of each strand[4,5]. The miRNA duplexes are 2’-O-methylated by the methyltransferase HEN1 at the 3′ terminus of each strand[6]. The duplex is loaded into an ARGONAUTE (AGO) protein to form a miRNA-induced silencing complex (miRISC). During miRISC formation, the duplex is unwound and one strand of the duplex is selected as the miRNA (or the guide strand), while the other strand called miRNA* (or the passenger strand) is ejected[7]. Most plant miRNAs are loaded into AGO1, which prefer miRNAs with a 5’ uridine (U)[8]. A miRISC binds to a target messenger RNA (mRNA) with sequence complementary to the miRNA, leading to the degradation or translational repression of the target transcript.

In plants, certain miRISCs, such as AGO1-miR173 and AGO7-miR390, can trigger the biogenesis of small interfering RNAs (siRNAs) from their target transcripts[9]. Upon cleavage of the target transcripts by these miRISCs, the cleaved products are converted to double-stranded RNAs (dsRNAs) by RNA-DEPENDENT RNA POLYMERASE 6 (RDR6) and the dsRNAs are processed by a Dicer protein, usually DCL4, to produce 21-nt siRNAs in a phased pattern[10,11]. Those siRNAs derived from the non-coding *TAS* loci are called trans-acting siRNAs (ta-siRNAs). In the Arabidopsis Columbia-0 (Col-0) genome, there are four families of *TAS* genes, *TAS1a/b/c, TAS2, TAS3a/b/c*, and *TAS4*[10,12–15]. The biogenesis of *TAS1a/b/c*- and *TAS2*-derived ta-siRNAs requires the AGO1-miR173 complex, and that of *TAS4*-derived ta-siRNAs requires the AGO1-miR828 complex[10,12,14]. However, for ta-siRNAs derived from the *TAS3a/b/c* loci, an AGO7-containing miRISC, AGO7-miR390, is required[16]. In addition to non-coding *TAS* loci, miRISCs such as AGO1-miR161.1 and AGO1-miR472 trigger the production of phased siRNAs (phasiRNAs) from protein-coding genes[17,18]. As in miRNA loading, ta-siRNA and phasiRNA duplexes are selectively loaded into AGO proteins based on their sequence features[8]. The ta-siRNA/phasiRNA-containing RISCs can direct target RNA cleavage and/or trigger the biogenesis of secondary siRNAs[9].

Eukaryotic AGO protein family members are highly conserved. An AGO protein contains four conserved domains: the N-terminal domain (N), which is required for small RNA duplex unwinding[19]; the PIWI/Argonaute/Zwille (PAZ) domain, which anchors the 3’ end of the miRNA guide strand[20,21]; the Middle (MID) domain, which binds the 5’ phosphate of the miRNA guide strand[22,23]; and the P-element-induced wimpy-testis (PIWI) domain, which in some AGOs, harbors an RNase H like motif that cleaves target RNA transcripts[24,25]. In Arabidopsis, AGO1, AGO2, AGO4, AGO7, and AGO10 have been shown to possess cleavage activity[7,13,26–28]. Structural studies of human AGO2 and prokaryotic AGOs show that the four domains of AGO form a two-lobed structure with a central cleft that cradles guide and target RNAs, with the N-PAZ domains composing one lobe and the MID-PIWI domains constituting the other[24,29]. The N and PAZ domain are directly connected by the L1 linker, while the MID and PIWI domain are directly connected by the L2 linker, which also connects the N-PAZ lobe and the MID-PIWI lobe[29–32].

In addition to the N, PAZ, MID and PIWI domains that are highly conserved, AGO proteins may harbor an N-terminal extension (NTE) of varying lengths and sequences, and the molecular or biological functions of this region are less understood. The NTE of AGOs is also referred as the N-terminal coil in some studies as it was predicted to possess coil-like structures. A recent study shows that a nuclear localization signal (NLS) and a nuclear export signal (NES) are present in the NTE (1-189a.a.) of Arabidopsis AGO1[33]. As inhibition of the EXPO1/NES-dependent protein nuclear export pathway significantly increases the ratio between nuclear and cytoplasmic AGO1, it was proposed that these signals direct the nucleo-cytosolic shuttling of AGO1[33]. Strikingly, AGO1 with the NES sequence mutated (AGO1mNES) associated with the same miRNA cohorts as its intact counterpart. This led to the proposal that the loading of miRNAs into AGO1 takes place in the nucleus and the NES mediates the nuclear export of miRISCs [33]. However, there are other nuclear export pathways that transport AGO1mNES from the nucleus to the cytoplasm. Indeed, the TREX-2 complex core component THP1 partners with the nucleoporin protein NUP1 at the nuclear envelope, together promoting the nuclear export of AGO1 or AGO1-miRISC[34]. Furthermore, the possibility of cytoplasmic AGO1 loading cannot be excluded. In fact, a cytoplasmically-sequestered AGO1 protein was shown to load miR165/6 *in vivo*[35].

56 mutant alleles of Arabidopsis AGO1 have been reported in the literature, with 20 of the 56 being missense mutations, however only one of the 20 alleles resides in the NTE[36]. *ago1-38*, a weak allele of *AGO1*, harbors a Gly-to-Arg mutation at the very end of the NTE region[37]. One study reported that the AGO1 protein abundance is similar between *ago1-38* and wild-type (WT), but the membrane association of AGO1 in the inflorescence tissue is reduced in *ago1-38*[38]. Other studies revealed that AGO1 and miRNAs are associated with membrane-bound polysomes, and the membrane-associated AGO1 could cause both target cleavage and translational repression[39,40]. Furthermore, *AGO1* is under feedback regulation - *AGO1* RNA is targeted by AGO1-bound miR168 with the miR168-binding site being located in the region encoding the NTE[41]. Studies in animal AGOs show diverse functions of the NTE region. In *Caenorhabditis elegans*, WAGO-1 (worm ARGONAUTE 1) and WAGO-3 are processed at their NTE by DPF-3, a dipeptidase[42]. Proteolytic activity of DPF-3 on the third and second a.a. of WAGO-1 and WAGO-3, respectively, promotes the correct sorting of 22G siRNAs into those AGOs and thus safeguard genome integrity[42]. *Drosophila melanogaster* AGO2 contains a ~400 a.a. long N-terminal glutamine-rich repeat (GRR) region, and reduced copy numbers of GRRs result in defects in RNAi responses and embryonic development[43].

Here, through analyzing the miRNA-related defects manifested by the Arabidopsis AGO1 NTE truncation lines, we uncover novel roles of the NTE in miRNA-mediated gene silencing. We show that loss of the entire NTE leads to a seedling-lethal phenotype resembling that of an *ago1* null allele, suggesting that the NTE is essential for the functions of AGO1. Additionally, we find that the 91-to-189a.a. region of the AGO1 NTE is essential for rescuing the morphological defects of the *ago1* null allele. Global analyses of miRNAs and their association with AGO1 revealed that the NTE, and particularly the 91-189 region, facilitates miRNAs’ loading into AGO1. Truncation of AGO1 1-to-90a.a. reduces its nuclear partitioning without affecting its profiles of miRNA and ta-siRNA association. Furthermore, we show that the biogenesis of ta-siRNAs requires the presence of either the 1-to-90a.a. or the 91-to-189a.a. region of AGO1. Taken together, our results reveal novel roles of the NTE of Arabidopsis AGO1.

## Results

### Truncation of the AGO1 N-terminal extension causes severe morphological defects

In *Arabidopsis thaliana*, the ten AGO family members can be grouped into three clades based on their phylogenetic relationship (Supplementary Fig.1a). All members of the AGO1/5/10 and AGO2/3/7 clades possess NTEs longer than 130 a.a., whereas members of the AGO4/6/8/9 clade contain NTEs less than 70 a.a. long (Supplementary Fig.1a). Protein sequence alignment of the NTE region of all Arabidopsis AGOs shows extremely low similarity (Supplementary Fig.1b). Even AGO10, the most closely related paralog of AGO1, shares little similarity in its NTE region to that of AGO1. Furthermore, phylogenetic analysis using only the NTE regions of the ten AGOs shows that the AGO10 NTE is the most distantly related to that of AGO1 (Supplementary Fig.1c). Together, the sequence analyses suggest that the NTE region of AGOs might serve AGO-specific functions.

To investigate the functions of the NTE (1-189a.a.) region of Arabidopsis AGO1, we searched for potential domains and functional motifs in this region by employing protein databases including Pfam, InterPro, and NCBI Conserved Domains Database. The 75-to-172 a.a region of Arabidopsis AGO1 was annotated as a glycine-rich_AGO1 domain by multiple protein databases. This domain appears in 117 species, all within the Magnoliopsida class (flowering plants), suggesting a unique function to flowering plants. Sequence alignment of the AGO1 NTE from six Magnoliopsida species, including *Arabidopsis thaliana (At), Arabidopsis lyrate (Al), Brassica napus (Bn), Glycine max (Gm), Oryza sativa (Os)*, and *Zea mays (Zm)*, shows that this glycine-rich_AGO1 domain is evolutionally conserved (Supplementary Fig.2). In addition, six RGG/RG repeats are found within the NTE region. The RGG/RG motif, which may undergo arginine methylation, is known for mediating RNA binding, protein localization, and protein-protein interactions [44]. Three of the six RGG/RG motifs are clustered at a.a. 83 to 103 in the glycine-rich_AGO1 domain, with two of them being conserved in AGO1s from the six species mentioned above except for *Oryza sativa*. One RGG/RG motif, ^59^RGGRG^63^, is conserved in all six species. Furthermore, glutamine (Q) content is particularly high in the NTE, with 30 Qs scattered along the region and accounting for 15.9% of the total amino acids in the NTE. This feature is conserved in the NTEs of all six inspected plant AGO1s. Next, we predicted sorting signals within the NTE that might facilitate the sub-cellular localization of AGO1. In addition to an NLS (^2^VRKRR^6^) and NES (^149^LAQQFEQLSV^158^) that were previously reported for Arabidopsis AGO1[33], a putative NLS (^102^GGGPSSGPPQ^111^) was predicted by SeqNLS with a score of 0.737 out of 1.

To understand how these features of the NTE affect the functions of AGO1, we tried to predict the structure of the NTE region by searching for similar protein sequences with available structural information in the Protein Data Bank using the SWISS-MODEL server. However, no protein template with significant similarity to AGO1 NTE was found. Next, we utilized AlphaFold, a machine learning tool that predicts protein structures, to get insights into the 3D structure of the NTE region (Fig.1a). The N-terminal segment (a.a 1-90) of the NTE is mostly unstructured and loosely attached to the rest of the AGO1 protein. The C-terminal segment (a.a. 91-189) of the NTE is more structured and could potentially interact with the PIWI domain of AGO1. The very C-terminus of the AGO1 NTE together with the L2 Linker is tucked in between the N and PIWI domains, and thus could potentially assist with the connection and movement between the N-PAZ lobe and the MID-PIWI lobe. The rest of the NTE C-terminus protrudes from the two lobes of AGO1, with a.a. 145-175 adopting an L shape that is close to the PIWI domain.

**Figure 1.**
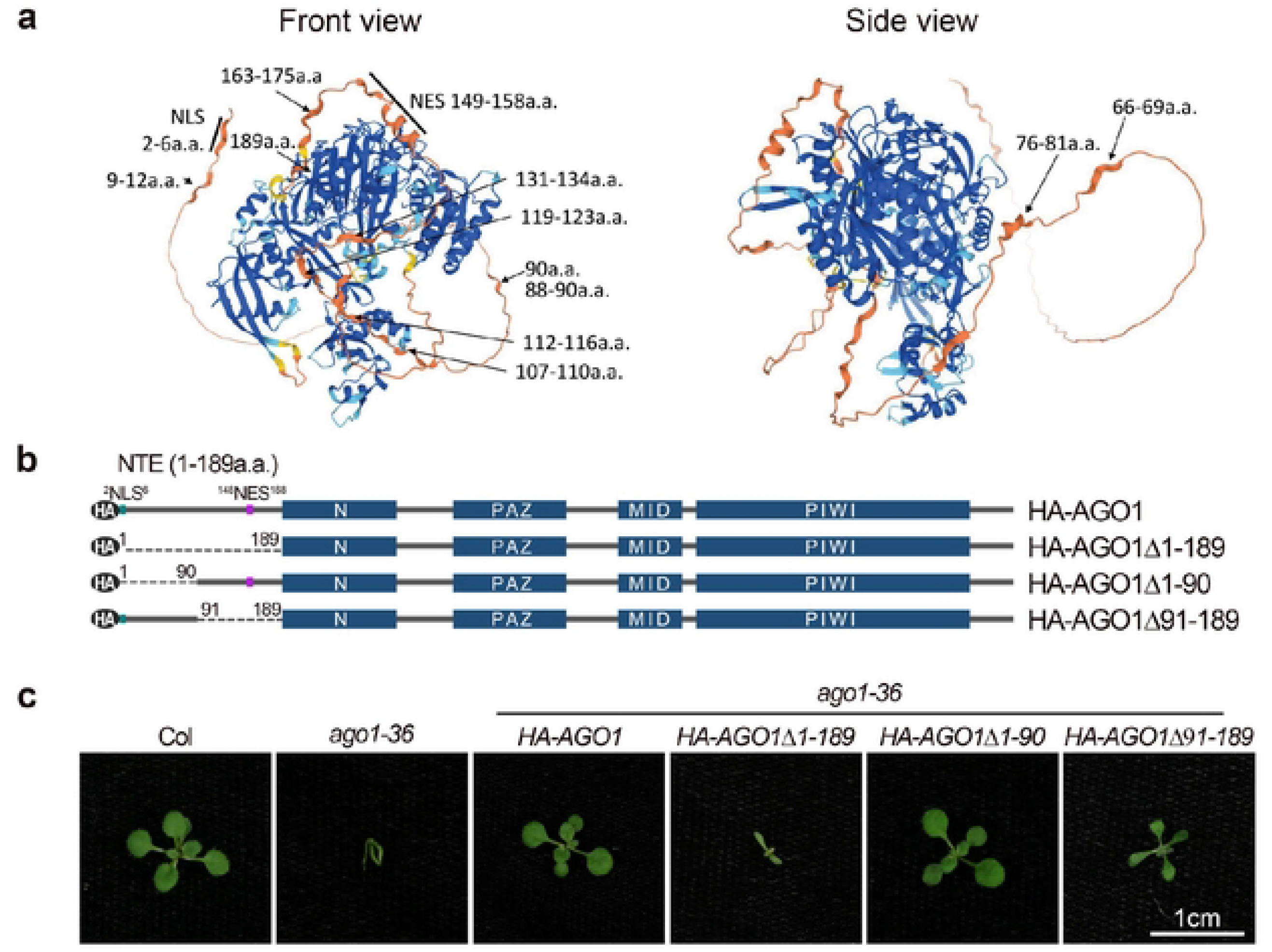
Arabidopsis AG01 harbors an unstructured N-terminal extension. (a) Front and side views of a putative Arabidopsis AG01 structure predicted by AlphaFold. The colors represent the per residue confidence score (pLDDT), with dark blue (pLDDT > 90), light blue (90 > pLDDT > 70), yellow (70 > pLDDT > 50), and orange (pLDDT < 50) denoting high, medium, low, and very low confidence, respectively. (b) Schematic representation of HA tagged Arabidopsis AG01 and AG01 NTE truncation forms. Blue blocks: domains; black ovals: the HA tag; solid lines: protein sequences; dashed lines: truncated protein sequences; green: NLS; purple: NE$. (c) Fourteen day-old plants of Col, *ago1-36,* and *ago1 36* expressing *H AAG01, HA-AG01Δ1-189, H A AG01Δ1-90,* and *HA AG01Ll91-189* transgenes. Scale bar, 1cm.

To further investigate how the NTE region affects the function of Arabidopsis AGO1 *in vivo*, we introduced N-terminal HA-tagged, wild-type (WT) or truncation mutants of AGO1 without the NTE region (AGO1Δ1-189), without the unstructured N-terminal segment of the NTE (AGO1Δ1-90), or without the more structured C-terminal segment of the NTE (AGO1Δ91-90) into the *ago1-36* mutant background (Fig.1b). The *ago1-36* allele contains a T-DNA insertion in the AGO1 gene at a position within the encoded PAZ domain[7]. The phenotype of *ago1-36* resembles *ago1* null alleles, such as *ago1-3*[45,46]. We found that both *HA-AGO1 ago1-36* and *HA-AGO1Δ1-90 ago1-36* had a WT phenotype, suggesting that the N-terminal half of the NTE is largely dispensable, at least under laboratory growth conditions. *HA-AGO1Δ91-90* partially rescued the developmental phenotypes of *ago1-36* plants, whereas *HA-AGO1Δ1-189 ago1-36* seedlings resembled *ago1-36* except that the hypocotyl hook was straightened, and the cotyledons were expanded at early developmental stages (Fig.1c). This suggests that the NTE, and particularly the region of a.a. 91-189, is essential for the functions of AGO1.

### The a.a. 91-189 region of the AGO1 NTE is crucial for miRNA accumulation

To determine the effects of AGO1 NTE truncation on miRNA accumulation, we performed RNA gel blot assays to determine the levels of miRNAs in seedling of *ago1-3, ago1-36,* and *ago1-36* expressing HA-tagged wild-type or truncated forms of AGO1. We found that compared to wild-type AGO1, the levels of three tested miRNAs were decreased in *AGO1Δ91-189* and *AGO1Δ1-189* such that they were similar to those of *ago1-36* and *ago1-3*, whereas their levels remained unchanged in *AGO1Δ1-90* (Fig.2a). Next, we used small RNA sequencing to examine the effects of AGO1 NTE truncations on miRNA accumulation. Principal component analysis (PCA) of the small RNA sequencing data showed that the three biological replicates of each genotype were highly reproducible (Supplementary Fig.3a). The size distribution of both total small RNAs and miRNAs was similar in *ago1-36* and *ago1-36* expressing wild-type or NTE-truncated AGO1s (Supplementary Fig.3b). Consistent with previous findings, total small RNAs showed a 21-nucleotide (nt) peak and a more abundant 24-nt peak, while the majority of miRNAs were 21-nt long[27,40].

**Figure 2.**
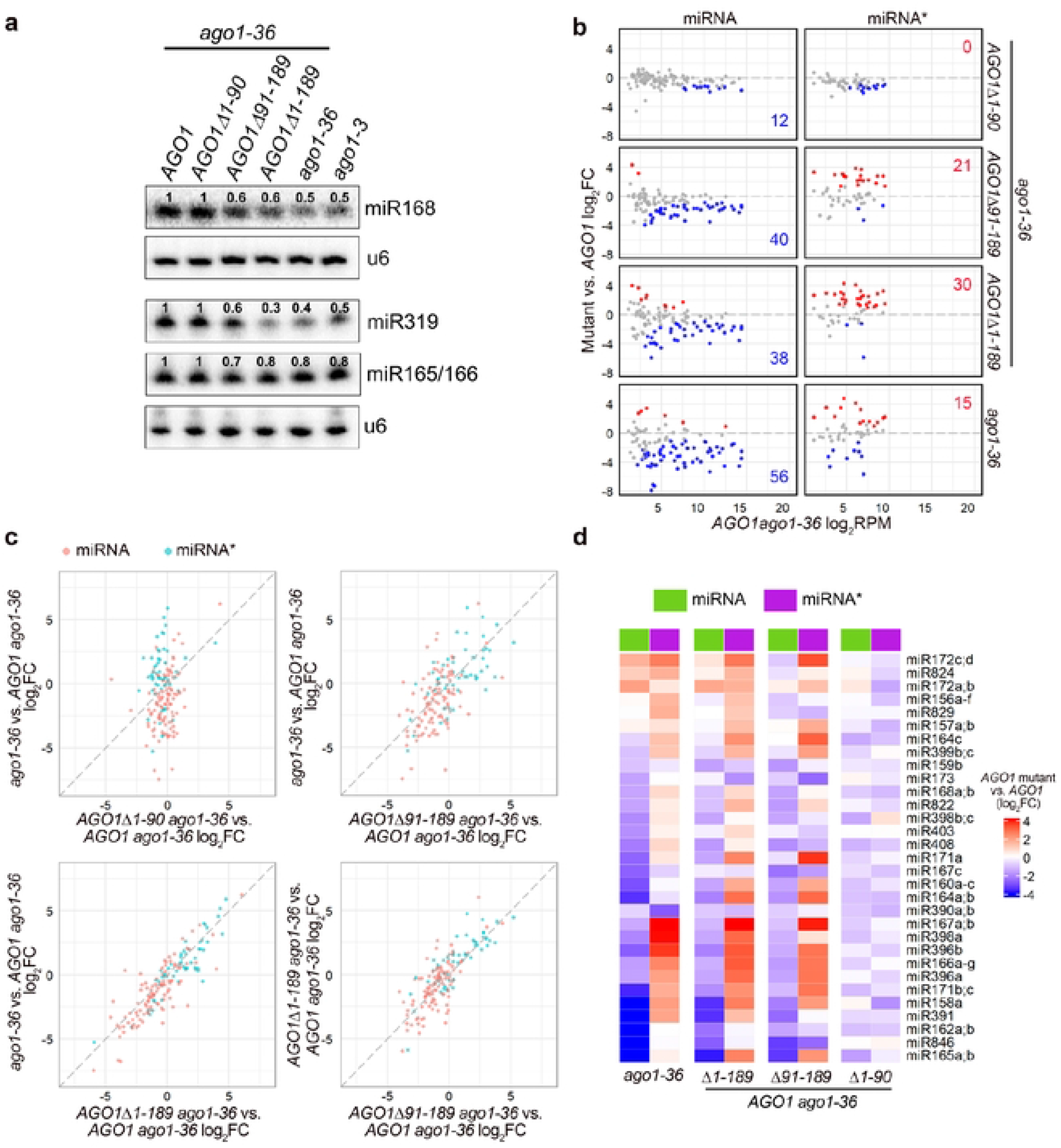
The AG01 N terminal extension is required for miRNA accumulation. (a) RNA gel blot analysis of miRNA abundance in 12-day-old seedlings of *ago1-3 ago1-36,* and *ago1-36* expressing *AG01* full-length or various NTE-truncated forms. The numbers represent miRNA abundance in different genotypes relative to *AG01 ago1 36.* The U6 blots serve as a loading control for the miRNA blots above. (b to d) Small RNA-seq analysis of 12-day-old seedlings. Except for *ago1-36,* all lines contain *pAG01:3XHA-AG01* or *pAG01:3XHA-AG01* NTE-truncated forms in the *ago1-36* background. Three biological replicates of each genotype were included in the analysis. (b) Scatter plots showing the abundance of miRNAs and miRNA*s in *AG01 ago136* (X axis) and fold change between *AG01* mutants and *AG01* (Y axis). Red and blue dots denote miRNAs or miRNA*s with significantly increased and decreased abundance (average RPM > 2, fold change > 1.5 and adjust *P* value < 0.01), respectively. (c) Scatter plots compar ng the log2(fold change) of miRNAs and miRNA*s between pairs of *AG01* mutants. (d) Heatmap depicting the levels of miRNAs and their corresponding miRNA*s in *AG01* mutants relative to *pAG01:3XHAAG01 ago1 36.*

Detailed analyses were carried out to compare the abundance of individual miRNAs between AGO1 NTE truncation lines. Among a total of 428 Arabidopsis miRNAs, 104 miRNAs and 53 miRNA*s with an average level > 2 RPM (reads per million) in the small RNA-seq samples were included in the analyses. Few differentially accumulated miRNAs and miRNA*s were found between *AGO1Δ1-90 ago1-36* and *AGO1 ago1-36*, with examples being miR165a/b, miR168a/b, miR171b/c and miR390a/b (Fig.2b and Supplementary Table 1). On the other hand, more than 36% of miRNAs were decreased and over 28% of miRNA*s were increased in *AGO1Δ91-189, AGO1Δ1-189*, and *ago1-36* when compared to *AGO1 ago1-36* (Fig.2b and Supplementary Table 1). This result was consistent with miRNA quantification using RNA gel blot assays (Fig.2a). In accordance with the previous report on *ago1-3*[46], a null allele of AGO1, around 54% (56 out of 104) of miRNAs were significantly decreased in *ago1-36* compared to *AGO1 ago1-36*; it is likely that failure of these miRNAs to load into an AGO protein made them more vulnerable to degradation. In contrast, 7 miRNAs were upregulated in *ago1-36* compared to *AGO1 ago1-36*, for example, miR172a;b, miR866, and miR5026. These upregulated miRNAs might be preferentially sorted into and stabilized by AGO2, as 5 out of 7 upregulated miRNAs possess a 5’ adenine (A), the preferred 5’ nucleotide by Arabidopsis AGO2[8]. In addition, 15 miRNA* accumulated to a higher level in *ago1-36* compared to *AGO1 ago1-36*, for example, miR391* and miR393a/b*, both of which containing a 5’ adenine (A) and shown to preferentially associate with AGO2[8]. As the majority of the upregulated miRNA*s in *ago1-36* contain a 5’ adenine (A) or guanine (G), they might be selectively loaded and protected by other AGOs when AGO1 is absent.

Comparison showed that the differentially accumulated miRNAs and miRNA*s (i.e. reduced levels of miRNAs and increased levels of miRNA*s) in *ago1-36* were highly correlated with those in *AGO1Δ1-189 ago1-36*, and with those in *AGO1Δ91-189 ago1-36*, although to a lesser extent (Fig.2c). No correlation was found for the differentially accumulated miRNAs and miRNA*s in *ago1-36* and *AGO1Δ1-90 ago1-36* (Fig.2c). These findings suggest that the AGO1 NTE (1-189a.a.), particularly the 91-189 region, is required for restoring miRNA and miRNA* levels in *ago1-36*. And as the differential accumulation of a small number of miRNAs and miRNA*s between *AGO1Δ1-189 ago1-36* and *AGO1Δ91-189 ago1-36* was not correlated, the a.a 1-90 region of the NTE might also play a minor role in miRNA accumulation.

We further assessed the effects of NTE truncations on miRISC formation by examining the abundance of miRNAs and their corresponding miRNA*s. During miRISC formation, AGO1 unwinds the miRNA/miRNA* duplex and selectively retains one strand (miRNA), whereas the other strand (miRNA*) is ejected[7]. A total of 31 miRNA species with both miRNA and miRNA* detected in the small RNA sequencing were evaluated. The levels of most miRNAs and their miRNA*s in *ago1-36* showed an inverse trend - while the majority of the miRNAs were downregulated, their corresponding miRNA*s were up-regulated, likely due to association with other AGOs (Fig.2d). Only a few miRNAs showed the same trend in changes in *ago1-36* to that of their corresponding miRNA*s, such as miR172a;b;c;d and miR390a;b (Fig.2d). A similar pattern was observed in *AGO1Δ1-189 ago1-36* and *AGO1Δ91-189 ago1-36*, indicating miRISC formation was likely defective for both forms of AGO1, and the a.a 91-189 region of AGO1 plays a major role in miRISC formation (Fig.2d).

### The a.a. 91-189 region of the AGO1 NTE is essential for miRNA and ta-siRNA loading

To test our hypothesis that the NTE region of AGO1 affects miRISC formation, we examined whether the association between miRNAs and ta-siRNAs with the NTE-truncated AGO1 is compromised. Small RNAs associated with the HA-tagged AGO1 and AGO1 mutants were immunoprecipitated (IP-ed) and sequenced. The AGO1Δ91-189 and AGO1Δ1-189 transgenes were in the *ago1-36* heterozygous background to ensure a similar cellular small RNA profile and morphological phenotypes across all tested plants. PCA analysis showed that the two biological replicates for each genotype were highly reproducible, and AGO1Δ91-189 IP and AGO1Δ1-189 IP samples were clustered together and separate from the other samples, suggesting they had small RNA profiles similar to each other yet different from AGO1 IP and AGO1Δ1-90 IP (Supplementary Fig.4a). Similar to total small RNA profiles (Supplementary Fig.3a), AGO1Δ1-90 IP clustered with AGO1 IP on the PC1 level, suggesting a similar small RNA binding preference between the two proteins (Supplementary Fig.4a).

Consistent with previous findings[8], the size distribution of wild-type AGO1 associated total small RNAs, miRNAs, and ta-siRNAs showed a 21-nt peak (Fig.3a). Like AGO1, AGO1Δ1-90 predominantly associated with 21-nt small RNAs, however, a minor reduced preference toward 21-nt miRNAs and an increased association with 20-nt miRNAs was observed (Fig.3a). As the sequenced size of small RNAs was used in the size distribution analysis, the 20-nt miRNAs could represent a pool of annotated 20-nt miRNAs, truncated miRNAs from a larger size, and misprocessed miRNAs. We specifically analyzed the association of miRNAs with various annotated sizes with AGO1. Annotated 20-nt miRNAs constituted a slightly higher proportion in AGO1Δ1-90-associated miRNAs as compared to wild-type AGO1, suggesting that the a.a. 1-90 region of the NTE might facilitate size selection of miRNAs (Supplementary Fig.4b). Strikingly, AGO1Δ91-189 and AGO1Δ1-189 completely lacked the preference toward 21-nt small RNAs, and their association with miRNAs and ta-siRNAs was minimal, suggesting that the a.a. 91-189 region of AGO1 is essential for miRNA and ta-siRNA loading (Fig.3a).

We next examined the differential association with individual miRNAs between AGO1 NTE mutants and wild-type AGO1. A total of 76 miRNAs and 24 miRNA*s were at an average level of > 2 RPM in the IP small RNA-seq samples and were included in the analysis. The association of miRNAs and miRNA*s with AGO1Δ1-90 was unaffected compared to wild-type AGO1, suggesting that a.a 1-90 of AGO1 have negligible effects on miRNAs loading. On the contrary, more than 88% of miRNAs and 87% of miRNA*s showed reduced association with both AGO1Δ91-189 and AGO1Δ1-189 (Fig.3b), and the miRNAs and miRNA*s downregulated in the two mutants were highly correlated (and Supplementary Fig.4c), suggesting that the 91-189 a.a. region of AGO1 affects miRNA association globally. Furthermore, both the miRNA strand and its corresponding miRNA* strand showed reduced association with AGO1Δ91-189 and AGO1Δ1-189 (Supplementary Fig.4d), implying that the reduced miRNA loading is unlikely due to defects in stand selection or miRNA* ejection. As AGO protein stabilizes its associated miRNAs, this compromised miRNA loading could explain the severe phenotypes and the reduced abundance of miRNAs in *AGO1Δ91-189 ago1-36* and *AGO1Δ1-189 ago1-36*. The reduced association of miRNA*s with AGO1Δ91-189 and AGO1Δ1-189 also suggests that the increased accumulation of miRNA*s in *AGO1Δ91-189 ago1-36* and *AGO1Δ1-189 ago1-36* is likely due to the loading of miRNA*s into other AGOs. Similar to miRNAs, most 21-nt ta-siRNAs showed reduced levels in AGO1Δ91-189 IP and AGO1Δ1-189 IP, but not in AGO1Δ1-90 IP (Fig.3c).

**Figure 3.**
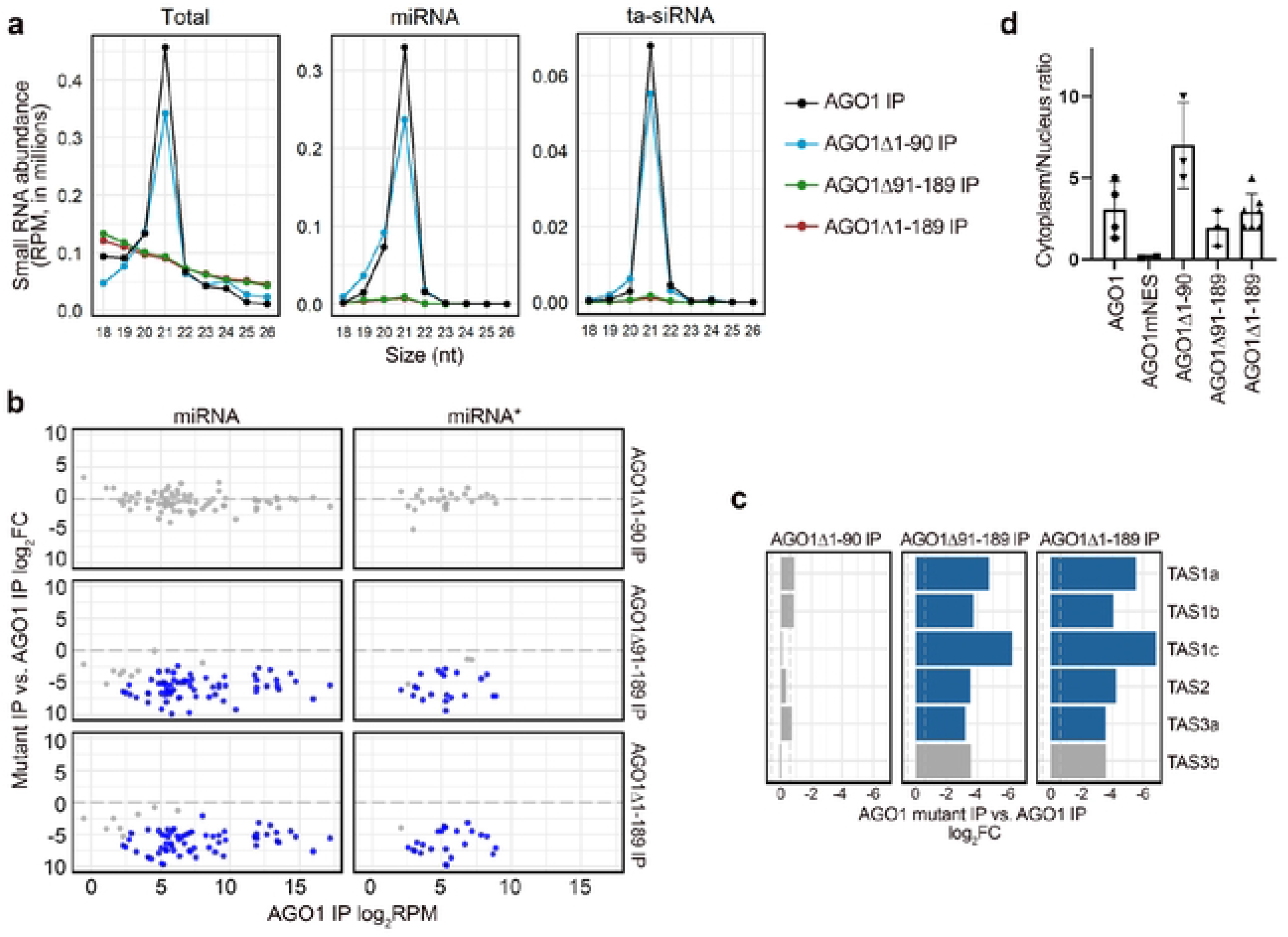
The loading of miRNAs and ta-siRNAs into AG01 is affected by truncations of the AG01 N terminal extension. (a to c) Analysis of small RNA seq data of AG01-associated small RNAs. Wild type AG01 and AG01Δ1-90 were in the *ago t36* background, while other AG01 NTE-truncated forms were in the *ago t36/+* background. Two biological replicates of each genotype were included in the analysis. (a) Size (in nucleotides (nt)) distribution depicting the abundance of 18 to 26 nt small RNAs in AG01 immunoprecipitates (IP) from plants containing AG01 and AG01 NTE-truncated forms. Total small RNAs, miRNAs and ta siRNAs are shown. RPM, reads per million (see Methods). (b) Scatter plots showing the miRNAs and miRNA*s with significantly different levels (fold change > 1.5 and adjust *P* value < 0.01) between IP-ed AG01 mutants and AG01. Red dots denote miRNAs or miRNA*s with significantly higher abundance, while blue dots denote miRNAs or miRNA*s with significantly lower abundance. (c) Bar plots depicting the log_2_(fold change) of 21-nt ta-siRNAs between IP-ed AG01 NTE-truncated forms and AG01. Blue bars denote ta-siRNAs with sign ficantly lower levels (fold change > 1.5 and adjust *P* value < 0.01). (d) Bar plots depicting the cytoplasmic/nuclear ratio of AG01 in AG01 NTE mutants. Wildtype AG01 and two replicates of AG01Δ190 were in the *ago 136* background, AG01mNES was in the Col background, while other AG01 NTEtruncated mutants and one replicate of AG01Δ1-90 were in the *ago1-36/+* background. Error bars indicate s.d.

### The 91-to-189 a.a region of AGO1 facilitates miRNA loading independently of its role in nuclear-cytoplasmic shuttling

AGO1 was previously reported to shuttle between the cytoplasm and the nucleus in an NLS (a.a. 2-6)- and NES (a.a. 149-158)- dependent manner[33]. The AGO1Δ1-90 protein lacks the NLS (a.a. 2-6), and the NES (a.a. 149-158) was removed from AGO1Δ91-189, while the AGO1Δ1-189 lacks both the NLS and the NES. To investigate whether the reduced loading of miRNAs and ta-siRNAs into AGO1 NTE mutants was due to defects in nuclear-cytoplasmic shuttling, we further examined the nucleocytoplasmic distribution of AGO1 in these mutants. In addition to the NTE truncation mutants generated in this study, we also included an NES mutated AGO1 (AGO1mNES) allele that was reported before, which showed enhanced nuclear localization, intact miRNA association, and reduced ta-siRNA binding[33]. Consistent with previous findings[33,34], the cytoplasm/nucleus ratio of wild-type AGO1 was around 3:1, and this ratio was reduced to 0.3:1 for the AGO1mNES protein (Fig.3d). In comparison to wild-type AGO1, the levels of cytoplasmic AGO1Δ1-90 increased, supporting the nucleus-importing role of NLS (a.a. 2-6) that resides within the 1-to-90 a.a. region of the NTE. However, the reduced levels of nuclear AGO1Δ1-90 did not affect its profiles of miRNA and ta-siRNA association (Fig.3b). Interestingly, the cytoplasm/nucleus ratio of the AGO1Δ1-189 protein lacking both NLS (a.a. 2-6) and NES (a.a. 149-158) was similar to that of wild-type AGO1, suggesting that there exist mechanisms to affect the nuclear-cytoplasm shuttling of AGO1 in an NTE-independent manner. Compared to wild-type AGO1, AGO1Δ91-189 showed increased nuclear accumulation, which is consistent with the role of the NES (a.a. 149-158) in directing AGO1’s nuclear export (Fig.3d). However, the truncation of a.a. 91-189 caused weaker effects in AGO1’s nuclear enrichment compared to the deletion of the NES. One possible explanation is that an unidentified motif in a.a. 91-189 of AGO1 facilitates the nuclear import. In this study, we identified a putative NLS (^102^GGGPSSGPPQ^111^) that resides in this region of AGO1, but its role in nuclear import has yet to be tested. It is worth noting that, unlike AGO1mNES, which predominantly resides in the nucleus and shows a wild-type-like miRNA-binding profile, miRNA association is depleted for AGO1Δ91-189 even though its nuclear accumulation is increased compared to wild-type AGO1, suggesting that a.a. 91-189 of AGO1 regulate miRNA loading independently of its role in the nuclear-cytoplasmic partitioning of AGO1 (Fig.3d).

### The 1-90 a.a. and 91-198 a.a. regions of the AGO1 NTE function redundantly in ta-siRNA biogenesis

To test whether the NTE region of AGO1 affects the activity of miRISC, we performed RNA-seq to examine the expression of miRNA target genes. The same RNA samples that were used in the small RNA sequencing analysis were also used here for the RNA-seq analysis, including *ago1-36* and *ago1-36* expressing wild-type *AGO1* or *AGO1* NTE truncated mutants driven by the *AGO1* promoter. Three biological replicates gave highly reproducible results (Supplementary Fig.5). 130 experimentally verified miRNA target genes were investigated to identify differentially expressed genes between wild-type *AGO1* and *AGO1* NTE mutants. In comparison to *AGO1 ago1-36*, gene expression in *AGO1Δ1-90 ago1-36* was largely unaffected (Fig.4a and 4b). Around 35% of miRNA target genes were derepressed in ago*1-36* and *AGO1Δ1-189 ago1-36* (Fig.4a), and the two genotypes were highly correlated in terms of the miRNA target genes affected (Fig.4c). Interestingly, only 25% of miRNA target genes were derepressed in *AGO1Δ91-189 ago1-36* (Fig.4a), and the levels of change were lesser than their counterparts in *ago1-36* and *AGO1Δ1-189 ago1-36* (Fig.4d). The less severe molecular phenotype of *AGO1Δ91-189 ago1-36* is consistent with their better seedling phenotype compared to *AGO1Δ1-189 ago1-36*, suggesting that a.a. 1-90 of AGO1 can partially rescue *AGO1Δ1-189 ago1-36*. Since miRNA association is reduced to a similar level in AGO1Δ91-189 and AGO1Δ1-189, it is possible that a.a. 1-90 of AGO1 facilitate target repression after miRNA loading.

**Figure 4.**
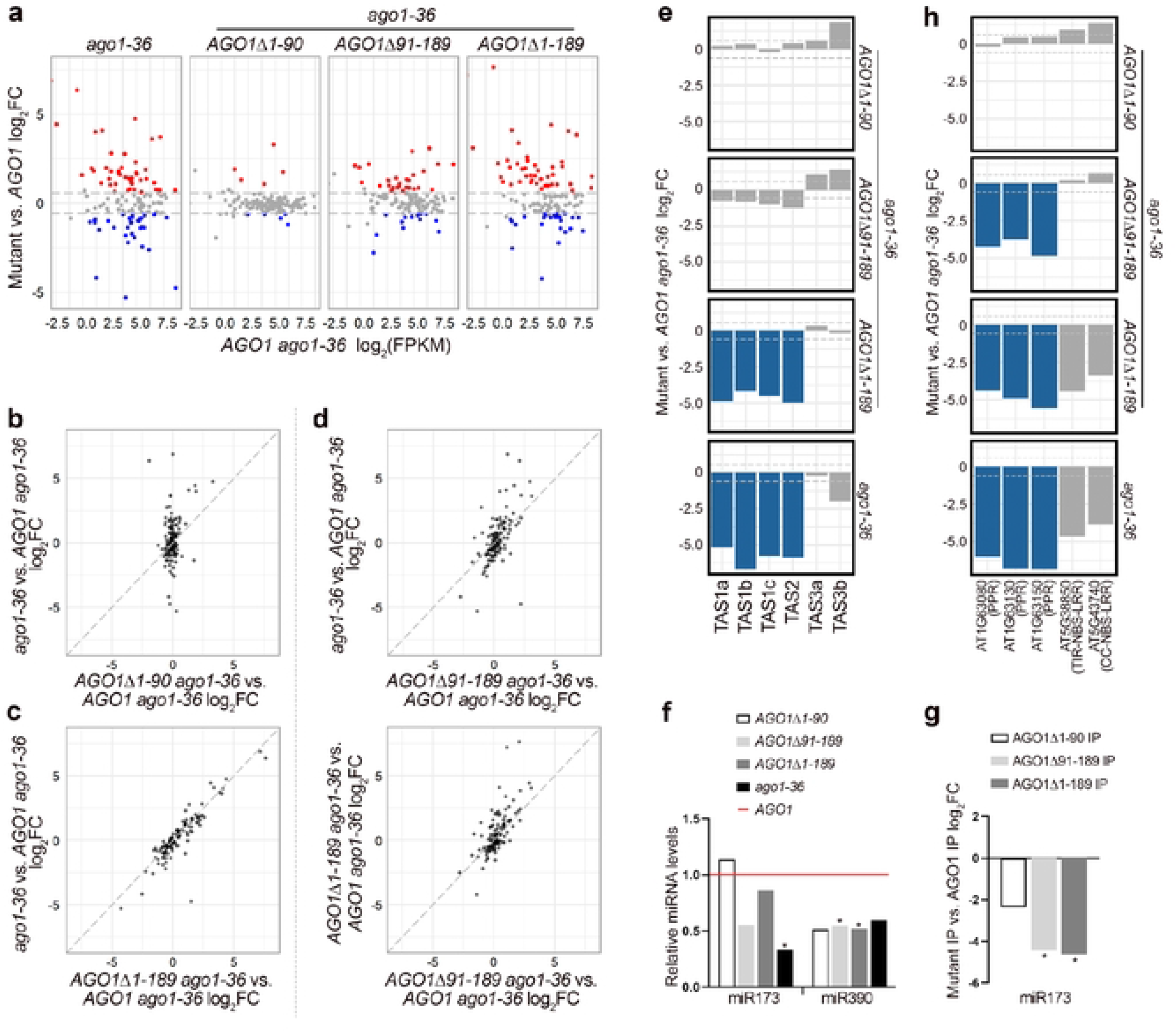
miRNA activities are affected by truncations of the AG01 N-terminal extension as revealed by RNA-seq and small RNA-seq analysis of 12-day-old seedlings. Wild-type AG01 and AG01 NTE truncated mutants were expressed in the *ago136* background. Three biological replicates of each genotype were included in the analysis. (a) Scatter plots showing miRNA target genes with significantly different expression (fold change > 1.5 and adjust *P* value < 0.05) between *AG01* mutants and *AG01.* Red and blue dots denote genes with sign ficantly increased and decreased expression, respectively. (b-d) Scatter plots comparing the log_2_(fold change) of miRNA target gene expression between pairs of *AG01* mutants. (e) Bar plots showing the log_2_(fold change) of 21-nt ta-siRNAs between *ago1-36* expressing various *AG01* mutants and *AG01 ago1-36* Blue bars denote ta-siRNAs with sign ficantly lower levels (fold change > 1.5 and adjust *P* value < 0.01). (f) Bar plots showing the relative levels of miR173 and miR390 in *ago136* expressing var ous *AG01* mutants compared to *AG01 ago1 36* (g) Bar plots showing the log_2_(fold change) of IP ed miR173 between AG01 NTE mutants and AG01. (h) Bar plots showing the log_2_(fold change) of 21-nt phasiRNAs between *ago1-36* expressing var ous *AG01* mutants and *AG01 ago1-36.* Blue bars denote phasiRNAs with significantly lower levels (fold change > 1.5 and adjust *P* value < 0.01).

In addition to target gene repression, certain miRNAs can trigger the biogenesis of secondary siRNAs from target loci such as *TAS1a/b/c* and *TAS2*, which produce ta-siRNAs in an AGO1-miR173 dependent manner. To determine whether the NTE of AGO1 affects ta-siRNA biogenesis, we compared the level of 21-nt small RNAs derived from the *TAS* loci and their phasing pattern. *TAS3a/b* ta-siRNAs, which require AGO7-miR390 for biogenesis, were included as a control. Strikingly, by small RNA-seq we found that ta-siRNA levels and their phasing pattern from the four AGO1-dependent loci were largely unchanged in *AGO1Δ1-90 ago1-36* or *AGO1Δ91-189 ago1-36* while the ta-siRNA levels were significantly reduced in *ago1-36* and *AGO1Δ1-189 ago1-36* (Fig.4e and Supplementary Fig. 6a). Consistently, RNA gel blot detection of *TAS1*-derived siR255 and *TAS2*-derived siR1511 showed unchanged levels in *AGO1Δ1-90 ago1-36,* and a moderate reduction in *AGO1ΔΔ1-189 ago1-36* while their levels were significantly reduced in *ago1-36* and *AGO1Δ1-189 ago1-36* (Supplementary Fig. 6b). The levels of *TAS3* ta-siRNAs were unaffected in *AGO1Δ1-189 ago1-36* (Fig.4e), consistent with expectations. We further confirmed that the reduced ta-siRNA levels from *TAS1* and *TAS2* loci in *AGO1Δ1-189 ago1-36* were not due to reduced accumulation of miR173, as the levels of miR173 were largely unaffected in all NTE truncation alleles of AGO1 (Fig.4f). Both AGO1Δ91-189 and AGO1Δ1-189 showed reduced association with miR173 (Fig.4g), which raised the possibility that miR173 triggered the biogenesis of ta-siRNAs in association with another AGO. However, this cannot explain the lack of ta-siRNA biogenesis in AGO1Δ1-189. A more likely scenario is that AGO1Δ91-189, despite reduced miR173 association, is sufficient to trigger ta-siRNAs production, while AGO1Δ1-189 cannot, which would suggest that a.a. 1-90 play a role in ta-siRNA biogenesis. However, ta-siRNA accumulation was not affected in *AGO1Δ1-90 ago1-36*, implying that a.a. 91-189 can also facilitate ta-siRNA biogenesis. It is likely that a.a. 1-90 and a.a. 91-189 of AGO1 redundantly enable the activities of AGO1 in ta-siRNA biogenesis.

Moreover, we noticed that phasiRNA production from protein-coding genes, such as *AGO1* and *PPR* genes, which are targeted by miR168 and miR161.1, respectively, was nearly abolished in *AGO1Δ91-189 ago1-36,* while phasiRNAs from *NBS-LRR* genes targeted by miR472 were largely unaffected (Fig.4h). However, the eliminated phasing pattern of *PPR*- and *NBS-LRR*-derived phasiRNAs in *AGO1Δ91-189 ago1-36* but not *AGO1Δ1-90 ago1-36* suggests that a.a. 91-189 are indispensable for miRNA-triggered phasiRNA biogenesis from protein-coding genes (Supplementary Fig. 6c). The absence of *AGO1*-derived phasiRNAs is due to the deletion of the miR168 targeting site within the a.a. 91-189 region of the NTE (Supplementary Fig. 6c). We further verified that the reduced phasiRNA levels from *PPR* and *NBS-LRR* genes in *AGO1ΔΔ1-189 ago1-36* were not due to reduced accumulation of their RNA transcripts (Supplementary Fig. 6d). Why is AGO1Δ91-189 capable of ta-siRNA biogenesis but not phasiRNA biogenesis from protein-coding genes? One possibility is that these two processes require different auxiliary proteins, and a.a. 91-189 are required for specific protein-protein interactions. Alternatively, these two processes might take place at distinct subcellular locations, and a.a. 91-189 are required for the localization of AGO1 to one of the subcellular compartments.

## Discussion

In this study, we show that the NTE (1-to-189 a.a.) region of AGO1 is essential for rescuing the developmental and molecular phenotypes of *ago1-36*. We further show that a.a. 91-189 are required for the association with miRNAs and ta-siRNAs *in vivo* independently of its role in nuclear-cytoplasmic shuttling and are thus essential for miRNA-mediated gene silencing. On the other hand, the a.a 1-90 region, which contains the NLS (a.a. 2-6), is dispensable for the association with miRNAs and ta-siRNAs *in vivo*. While the NTE is essential for ta-siRNA biogenesis, the presence of either a.a. 1-90 or a.a. 91-189 of the NTE is sufficient for ta-siRNA biogenesis, suggesting a redundant function of these two regions in ta-siRNA biogenesis.

Proper loading into an AGO protein is essential for miRNAs to exert their roles in target gene repression. We show that miRNA association is severely and similarly compromised for AGO1Δ91-189 and AGO1Δ1-189 (Fig.3a, 3b, and Supplementary Fig.4c), suggesting a.a. 91-189 of AGO1 are required for miRNA loading. Whereas AGO1Δ1-90 IP shows a similar miRNA profile to wild-type AGO1 IP, suggesting that a.a. 1-90 of AGO1 are not required for miRNA loading (Fig.3a and 3b). A previous study shows that an NLS (a.a. 2-6) in AGO1 promotes the nuclear localization of a reporter protein (GFP-GUS) when a section of the AGO1 NTE (a.a. 1-148) was fused to the reporter protein, while mutating an NES (a.a. 149-158) prevents the nuclear export of AGO1[33]. A similar finding was made in this study, we show that compared to wild-type AGO1, the cytoplasmic/nuclear ratio of AGO1Δ1-90 and AGO1Δ91-189 was increased and decreased, respectively (Fig 3d). It was proposed that miRNA loading into AGO1 occurs inside the nucleus, as AGO1mNES, which predominantly resides in the nucleus, shows a similar miRNA binding profile to that of wild-type AGO1[33]. In this study we show that AGO1Δ91-189, which exhibits increased nuclear localization, does not manifest a wild-type-like miRNA-binding profile as does AGO1mNES, suggesting that a.a. 91-189 of AGO1 are critical to miRNA loading independently of its role in the nuclear-cytoplasmic shuttling of the protein (Fig 3b and 3d). It is possible that a.a. 91-189 of AGO1 are required for the interaction with HSP90, which is essential for miRNA loading[47]. Furthermore, we show that the reduced nuclear localization of AGO1Δ1-90 does not affect its miRNA loading or target transcripts repression, suggesting that the residual nuclear localization was sufficient to allow for miRNA loading or that miRNA loading can also occur in the cytoplasm (Fig 3b, 3d and 4a). To test whether miRNA loading can occur in the cytoplasm, an AGO1 mutant exclusively residing in the cytoplasm is required. Unlike AGO1mNES, which almost exclusively resides in the nucleus, 30% of the AGO1Δ91-189 protein is in the cytoplasm, suggesting a motif that promotes nuclear import could be in this region (Fig 3d). An NLS (^102^GGGPSSGPPQ^111^) that resides in the a.a. 91-189 region of AGO1 was predicted in this study. Mutating both NLS (a.a. 2-6) and NLS (a.a. 102-111) might create a cytoplasm-exclusive AGO1 that is suitable to test our hypothesis.

The 1-to-90 a.a. region of AGO1 is not required for rescuing the developmental defects of *ago1-36* or for miRNA loading, however, we found it can facilitate target repression and ta-siRNA biogenesis. We show that the association of miRNAs with AGO1Δ91-189 and AGO1Δ1-189 is almost abolished (Fig.3a and 3b), whereas miRNA target gene repression is less affected in *AGO1Δ91-189 ago1-36* compared to *AGO1Δ1-189 ago1-36* (Fig 4a and 4d), suggesting that a.a. 1-90 of AGO1 assist in target transcript cleavage. Furthermore, although miR173, the trigger miRNA for the biogenesis of *TAS1a/b/c*- and *TAS2*-derived ta-siRNAs, was significantly reduced in AGO1Δ91-189 IP compared to wild-type AGO1 IP (Fig 4g), ta-siRNA biogenesis was largely unaffected in *AGO1Δ91-189 ago1-36* (Fig 4e). It is unlikely that other AGOs associate with miR173 to trigger ta-siRNA biogenesis when AGO1 loading is compromised, as miR173-dependent ta-siRNA biogenesis is downregulated in *ago1-36* (Fig 4e). Together, these observations suggest that a.a. 1-90 of the NTE are sufficient for ta-siRNA biogenesis. We hypothesize that only a small amount of AGO1-miR173 is required to trigger the biogenesis of ta-siRNAs.

Although miRNAs and ta-siRNAs were almost depleted in AGO1Δ91-189 IP and AGO1Δ1-189 IP, these proteins are still associated with small RNAs *in vivo* (Fig.3a). Wild-type AGO1 is mainly associated with 21-nt miRNAs and ta-siRNAs, although it is also loaded with Pol IV-dependent siRNAs and small RNAs derived from coding genes, TEs, inverted-repeats, snRNA, snoRNA, rRNA, and tRNA. We found that over 80% of the 18-to-26 nt reads from the small RNA libraries from AGO1Δ91-189 and AGO1Δ1-189 IP are derived from rRNA (Fig.5a). The proportion of rRNA- and tRNA-derived small RNAs are dramatically increased in AGO1Δ91-189 IP and AGO1Δ1-189 IP compared to wild-type AGO1 IP, while all other small RNA species show a corresponding decrease (Fig.5a). We suspect that when lacking a.a. 91-189, AGO1 is unable to distinguish Dicer-dependent small RNAs from rRNA-derived small RNAs. This would be an exciting avenue for future investigation.

**Figure 5.**
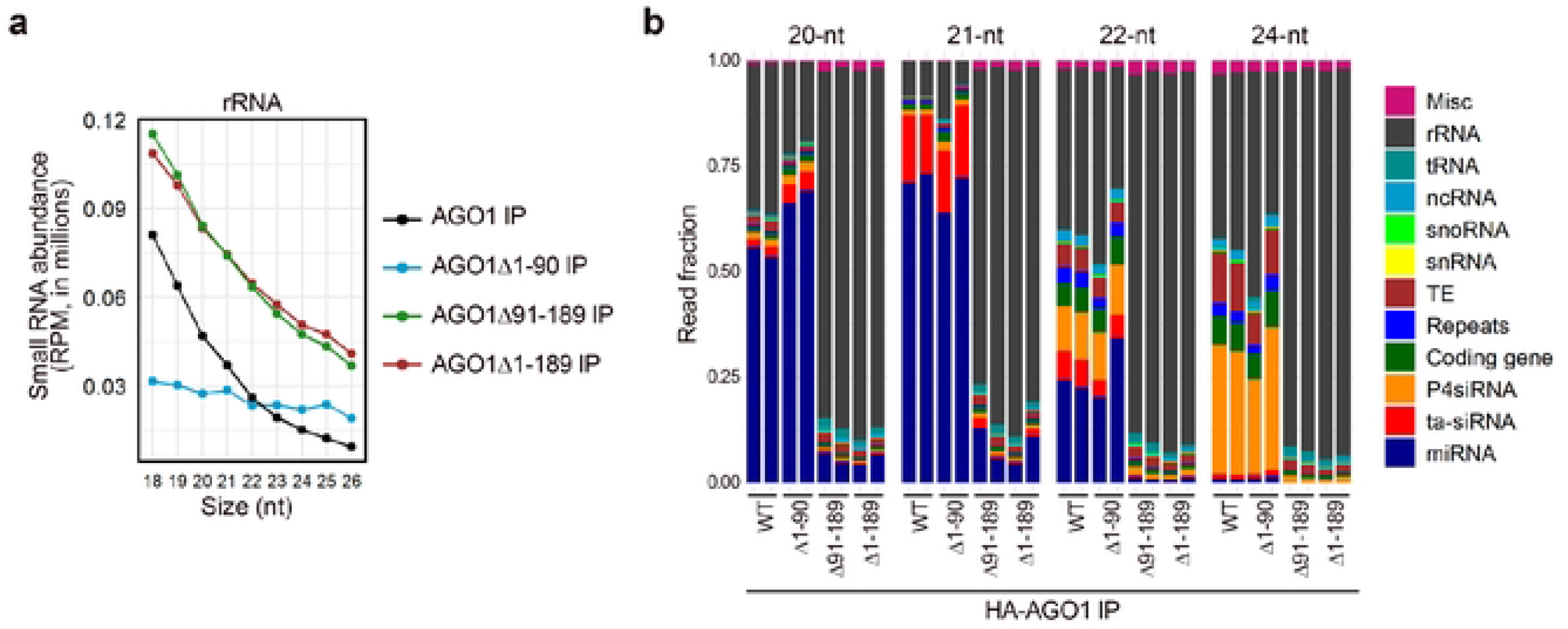
The N-terminal extension of AG01 allows AG01 to distinguish between Dicer­ dependent small RNAs and rRNA-derived small RNAs (rsRNAs). (a to b) Analysis of small RNA seq data of AG0 1associated small RNAs. Wildtype AG01 and AG01Δ1-90 were in the *ago t36* background, while other AG01 NTE­ truncated mutants were in the *ago 136/+* background. Two biological repl cates of each genotype were included in the analysis. (a) Size (in nucleotides (nt)) distribution depicting the abundance of 18- to 26-nt rsRNAs in IPs from AG01 and AG01 NTE-truncated mutants. RPM, reads per million (see Methods). (b) Composition of small RNAs in the 20-, 21-, 22-, and 24 nt classes in IPs from wild­ type AG01 (WT) and AG01 NTE truncated forms. Each column represents a biological replicate.

## Methods

### Plant materials and growth conditions

All *Arabidopsis thaliana* lines in this study were in the Columbia (Col) ecotype. Seeds of *ago1-36* (SALK_087076) were obtained from the ABRC collection. Seeds were surface-sterilized with 75% ethanol and stratified in water at 4°C for 2 days then transferred to 1x Murashige–Skoog (MS) medium. All plants were kept in a growth chamber at 22°C under full-spectrum white light and long-day conditions (16 h light / 8 h dark).

*ago1-36* and *ago1-36/+* expressing *pAGO1∷3XHA-AGO1*, *pAGO1∷3XHA-AGO1Δ1-90, pAGO1∷3XHA-AGO1Δ91-189,* and *pAGO1∷3XHA-AGO1Δ1-189* were generated in this study. To construct the plasmids of the above transgenes, a 1648bp *AGO1* promoter (including the 5’ UTR) and the 399bp 3’ UTR were amplified from genomic DNA, the full length *AGO1* coding region and NTE-truncated *AGO1* were amplified from cDNA, and the 3xHA tag was amplified from pGWB615[48]; these fragments were then cloned into pEarleyGate 301[49] (predigested by XbaI and NcoI) using the NEBuilder HiFi DNA Assembly Cloning Kit (NEB). All clones were validated by sequencing. Sequences of the primers are listed in Supplementary Table 2.

### Small RNA gel blot analysis and sequencing

RNA isolation and RNA gel blot analysis of small RNAs were performed as described[40]. 5-10 μg of total RNA was isolated from 12-day-old Arabidopsis seedlings using TRI Reagent (NRC), resolved in 15% urea–PAGE gels and transferred to NX membranes (Amersham Hybond-NX). MiRNAs were detected with 5’ end ^32^P-labeled antisense DNA oligonucleotide probes. Sequences of the DNA oligonucleotide probes in this study are listed in Supplementary Table 2. Hybridization signals were analyzed by phosphorimager (Typhoon 9410, GE).

Small RNA sequencing and data analysis were performed as described[40]. 25 μg of total RNA was extracted from 12-day-old Arabidopsis seedlings and resolved in a 15% urea-PAGE gel, and small RNAs of 15- to 40-nt were isolated from the gel. Small RNA libraries of gel-purified small RNAs or small RNAs acquired from AGO1 immunoprecipitation (as described below) were constructed using the NEBNext Multiplex Small-RNA Library Prep Set for Illumina (E7300). The libraries were sequenced on an Illumina HiSeq X Ten, and the resulting data were analyzed using an in-house pipeline pRNASeqTools v.0.8 (https://github.com/grubbybio/pRNASeqTools). The three-prime-end adapter sequence (AGATCGGAAGAGC) was trimmed from raw reads, followed by filtering to retain 18- to 42-nt reads using cutadapt v3.0[50]. Trimmed reads were then aligned to the Arabidopsis genome (TAIR10) using ShortStack v.3.8.5[51] with parameters ‘-bowtie_m1000 -ranmax 50 -mmap u -mismatches 0’. For total small RNAs, reads were normalized against total 18- to 42-nt reads minus 45S rRNA reads (RPM, Reads Per Million Reads). For small RNAs acquired from AGO1 immunoprecipitation, reads were normalized against IP-ed 18- to 42-nt reads (RPM). Differential comparison of small RNAs was conducted by DESeq2 v1.30.0 with fold change of 1.5 and adjusted *P* value < 0.01 as the parameters[52]. Annotation of miRNA and miRNA* sequences was obtained from miRbase v21 (http://www.mirbase.org/). For miRNA length distribution and small RNA composition analysis, reads that match perfectly or with a 1-nt shift on either end from the annotated sequence were assigned to the miRNA. For miRNA differential expression analysis, only reads that match perfectly to the annotated sequence were assigned to the corresponding miRNA. For ta-siRNA, levels were quantified by summing 21-nt small RNA reads that mapped to each of the eight *TAS* loci.

### Immunoprecipitation and protein gel blot analysis

2 g of 12-day-old seedlings was ground into fine powder in liquid nitrogen and the powder was homogenized in 3 ml lysis buffer (50 mM Tris-HCl, pH7.5, 150 mM NaCl, 10% glycerol, 0.1% CA-630, one tablet of cOmplete EDTA-free Protease Inhibitor Cocktail /50ml (Roche)), and incubated for 30 min at 4°C with gentle rotation. The total lysate was centrifuged at 12,000 g at 4°C for 20 min twice to remove cell debris. Meanwhile, 30 μl Dynabeads™ Protein A (ThermoFisher) was incubated with 8 μg of anti-HA antibody (Sigma, H6908) for 30 min at room temperature with gentle rotation, and the beads were then washed with lysis buffer 5 times to remove the extra antibody. The lysate was incubated with the anti-HA antibody-protein A beads for 2 hours at 4°C with gentle rotation, and then beads were captured magnetically and washed with lysis buffer 5 times. Washed beads were divided for small RNA and protein analysis.

For small RNA analysis, washed beads were boiled in H_2_O for 5 min under constant shaking and removed magnetically. The supernatant containing immunoprecipitated RNA was used in small RNA library construction (as described above). For protein gel blot analysis, washed beads were boiled in 2xSDS sample buffer (50 mM Tris-HCL at pH 6.8, 10% glycerol, 2% SDS, 0.1% bromophenol blue, and 1% 2-mercaptoethanol) for 10 min followed by vigorous shaking. The beads were then removed magnetically, and the supernatant containing the immunoprecipitated protein was resolved in a 10% SDS-PAGE gel and detected by an anti-HA antibody (Roche, 12158167001).

### RNA-seq and data analysis

Total RNA was isolated from 12-day-old Arabidopsis seedlings using TRI Reagent (NRC), and DNA was removed by DNase I (Roche) treatment. PolyA RNA was then enriched from 1 μg of DNase I-treated RNA using Oligo d(T)25 Magnetic Beads (NEB S1419S), followed by RNA-seq libraries construction using the NEBNext Ultra Directional RNA Library Prep Kit for Illumina (NEB E7420). RNA libraries were sequenced on an Illumina NovaSeq 6000 platform (PE150 bp), and the sequencing data were analyzed using the pRNASeqTools v.0.8 pipeline. Briefly, raw reads were aligned to the Arabidopsis genome (TAIR 10) using STAR v2.7.6a[53] with parameters ‘--alignIntronMax 5000 --outSAMmultNmax 1 --outFilterMultimapNmax 50 --outFilterMismatchNoverLmax 0.1’, and counted by featureCounts v2.0.0[54]. Normalization was performed by calculating the FPKM (Fragments Per Kilobase Million) for each gene, and differential gene expression analysis was conducted by DESeq2 v1.30.0 with a fold change of 1.5 and adjusted *P* value < 0.05 as the parameters[52].

### Small RNA phasing analysis

Small RNA phasing analysis was performed as previously described[18,40,55]. Small RNA reads from both the sense and antisense strands were included in the analysis. The formula used for phasing score calculation was described before[40].

### Nuclear–cytoplasmic fractionation

Nuclear-cytoplasmic fractionation was performed as described[34]. 1 g of 12-day-old Arabidopsis seedlings was crosslinked in 0.5% formaldehyde/1× PBS buffer through vacuum infiltration for 15 min at room temperature, and crosslinking was stopped by vacuum infiltration in 100 mM glycine/1× PBS buffer for 5 min at room temperature. The plant material was washed with 1× PBS buffer and blotted dry, and then ground to a fine powder in liquid nitrogen. The powder was resuspended in 2 ml lysis buffer (20 mM Tris-HCl, pH7.5, 20 mM KCl, 2.5 mM MgCl_2_, 2 mM EDTA, 25% glycerol, 250 mM sucrose, 5 mM DTT and cOmplete Protease Inhibitor Cocktail (Roche)) and filtered through a 40 μm cell strainer (Falcon). The flow-through was centrifuged at 1,500g for 10 min at 4°C. The supernatant representing the cytoplasmic fraction was further centrifuged at 10,000g for 10 min at 4°C to remove any residual debris. The pellet from the 1,500g spin representing the nuclei was dissolved with 10 ml nuclei resuspension buffer 1 (NRB1) (20 mM Tris-HCl, pH7.4, 2.5 mM MgCl_2_, and 0.2% Triton X-100) and centrifuged at 1,500g for 10 min to collect nuclei; this step was repeated 8 times to thoroughly wash the nuclei. After the final wash, the pellet was resuspended with 500 μl NRB2 (20 mM Tris-HCl, pH7.5, 250 mM sucrose, 10 mM MgCl_2_, 0.5% Triton X-100, 5 mM 2-mercaptoethanol, and cOmplete Protease Inhibitor Cocktail (Roche)), and carefully loaded onto 500 μl NRB3 (20 mM Tris-HCl, pH7.5, 1.7 M sucrose, 10 mM MgCl_2_, 0.5% Triton X-100, 5 mM 2-mercaptoethanol, and cOmplete Protease Inhibitor Cocktail (Roche)) without disturbing the bottom layer, then the sample was centrifuged at 16,000g for 45 min at 4°C. The nuclear pellet was resuspended and boiled in 1XSDS loading buffer for 10 min for protein gel blot analysis.

### Multiple sequence alignment

Protein sequences of AGO1 paralogs and orthologs from *Arabidopsis thaliana* and representative species were acquired from Uniprot (https://www.uniprot.org/), including 10 *Arabidopsis thaliana* AGOs, AGO1 (O04379), AGO2 (Q9SHF3), AGO3 (Q9SHF2), AGO4 (Q9ZVD5), AGO5 (Q9SJK3), AGO6 (O48771), AGO7 (Q9C793), AGO8 (Q3E984), AGO9 (Q84VQ0), and AGO10 (Q9XGW1), *Arabidopsis lyrata* AGO1 (D7KD09), *Brassica napus* AGO1 (A0A078JMZ3), *Glycine max* AGO1a (I1MQL3), *Oryza sativa* AGO1a (Q6EU14), and *Zea mays* AGO1a (A0A096TTL7). Protein sequence alignments were conducted by MUSCLE [56] (https://www.ebi.ac.uk/Tools/msa/muscle/) with default settings, and figures were made using ESPript3[57] (https://espript.ibcp.fr).

### Structure prediction

The protein sequence of *Arabidopsis thaliana* AGO1 (O04379) was obtained from Uniprot (https://www.uniprot.org/). Structure prediction was conducted by AlphaFold[58] (https://alphafold.ebi.ac.uk/) with default settings. Protein structure homology-modelling was performed by SWISS-MODEL (https://swissmodel.expasy.org/).

### Protein feature analysis

Protein domain and motif analysis was performed by Pfam (http://pfam.xfam.org/), InterPro (https://www.ebi.ac.uk/interpro/), and NCBI Conserved Domains Database (https://www.ncbi.nlm.nih.gov/Structure/cdd/wrpsb.cgi. Protein subcellular localization prediction was performed by WoLFPSORT (https://wolfpsort.hgc.jp/), SeqNLS (http://mleg.cse.sc.edu/seqNLS/), cNLS Mapper (http://nls-mapper.iab.keio.ac.jp/cgi-bin/NLS_Mapper_form.cgi), and NESmapper (https://sourceforge.net/projects/nesmapper/).

## Data availability

Raw sequence data and processed data generated in this study were deposited in the NCBI GEO database. The small RNA-seq and mRNA-seq data of *ago1-36* and *HA-AGO1ago1-36* are under the accession number GSE176568. And the small RNA-seq and mRNA-seq data of AGO1 NTE mutants are under the accession number GSE212365. Other supporting data is available from the corresponding author upon request.

## Author Contributions

Y.X. and X.C. designed the project; Y.X. generated and characterized the transgenic lines, prepared sequencing libraries, performed bioinformatic analysis, and conducted *At*AGO1 structure prediction and AGO sequence alignments; Y.Z., Z.F.L., A.S., and S.P. screened the transgenic lines; Y.X. and X.C. analyzed data; Y.X. prepared the figures; Y.X. and X.C. wrote the manuscript and all authors revised the manuscript.

## Acknowledgments

We would like to thank Dr. Nicolas Bologna for sharing the *pAGO1∷GFP-AGO1mNES(WT)* transgenic line. We thank Chenjiang You and Brandon H. Le for advice in bioinformatics data analysis, and Yuan Wang for discussions. This work was supported by funds from NIH GM129373 to X.C.

## Supplementary Figures and Figure Legends

**Figure S 1.**
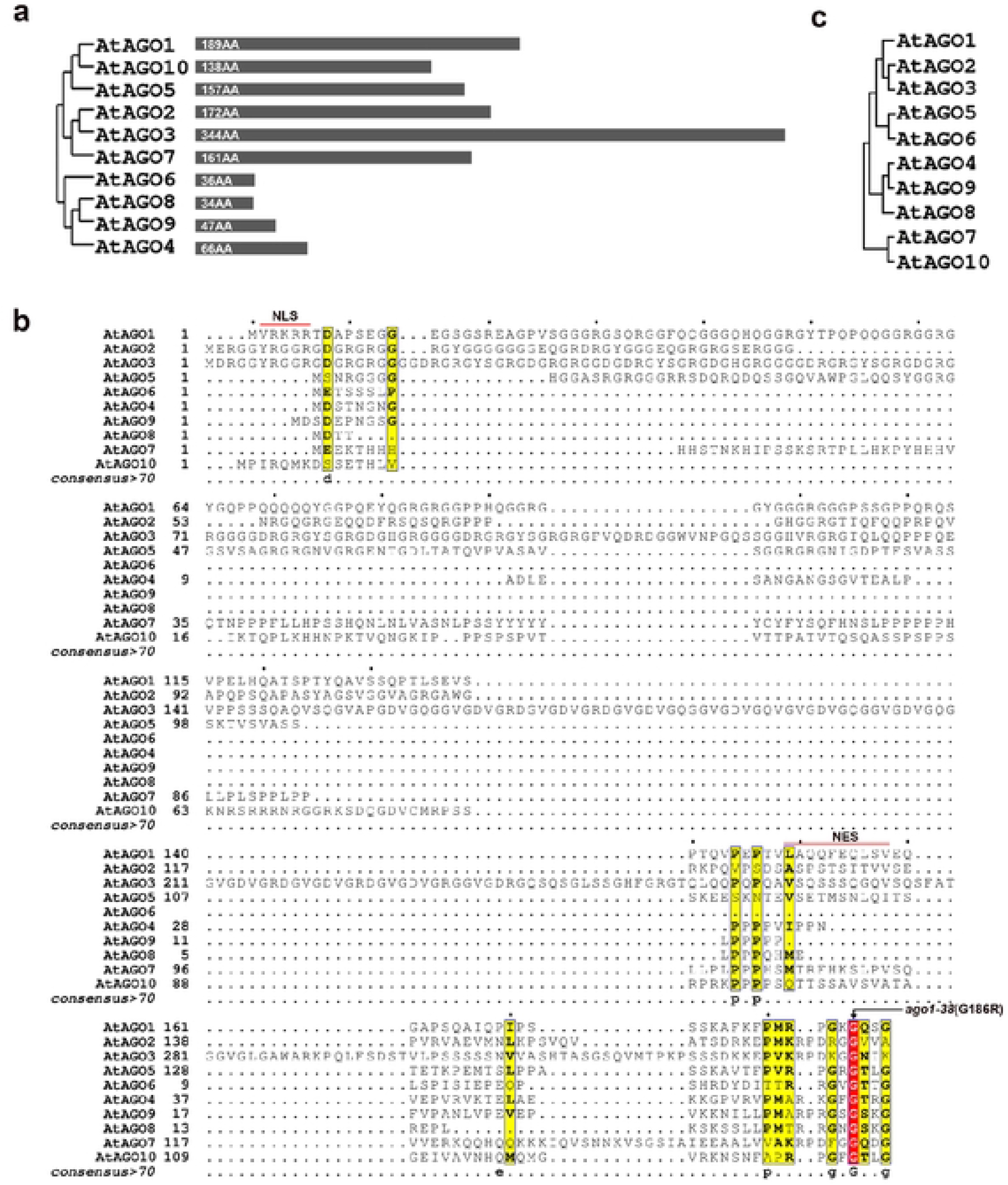
Phylogenetic analysis ofArabidopsis AGOs and Arabidopsis AGO NTEs, and sequence alignment of Arabidopsis AGO NTEs. (a) Phylogenetic analysis of 10 Arabidopsis AGOs. The black bars denote the lengths of the NTE regions in the AGOs. (b) Alignments showing the conservation or lack of conservation of the NTE regions in the Arabidopsis AGO family. The red rectangle denotes identity of residues in all proteins. Yellow rectangles denote similar residues. The NLS and NES are marked by the red lines. (c) Phylogenet c analysis of the NTE regions of 10 Arabidopsis AGOs.

**Figure S2.**
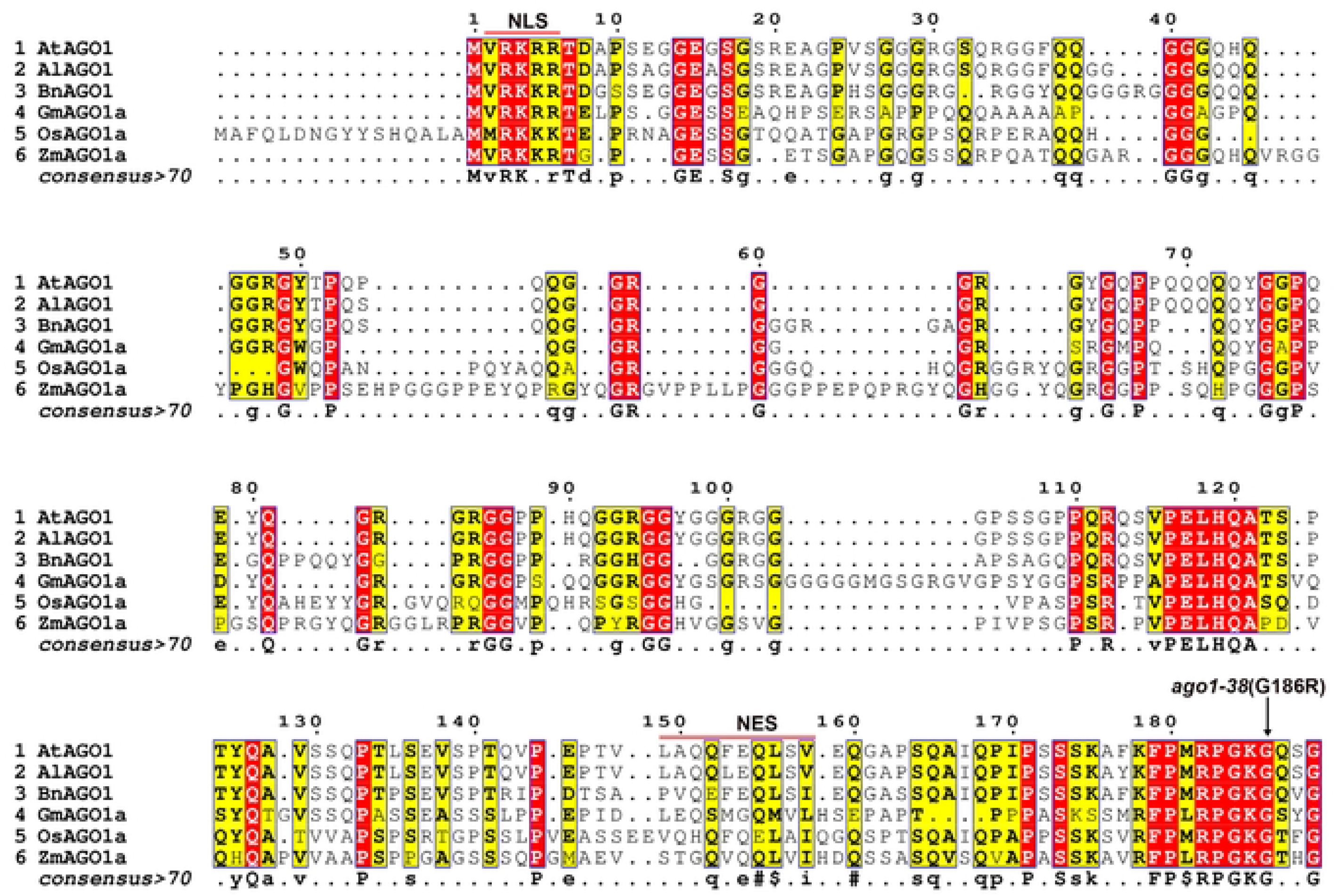
Sequence alignments showing the conservation or lack of conservation of *Arabidopsis thaliana* AG01 NTE regions in AG01 s from six plant species, including *Arabidopsis thaliana (At), Arabidopsis lyrate (Al), Brassica napus (Bn), Glycine max (Gm), Oryza sativa (Os)*, and *Zea mays (Zm)*. The red rectangles denote amino acid identity in all proteins. Yellow rectangles denote similar residues. The NLS and NES are marked by the red lines.

**Figure S3.**
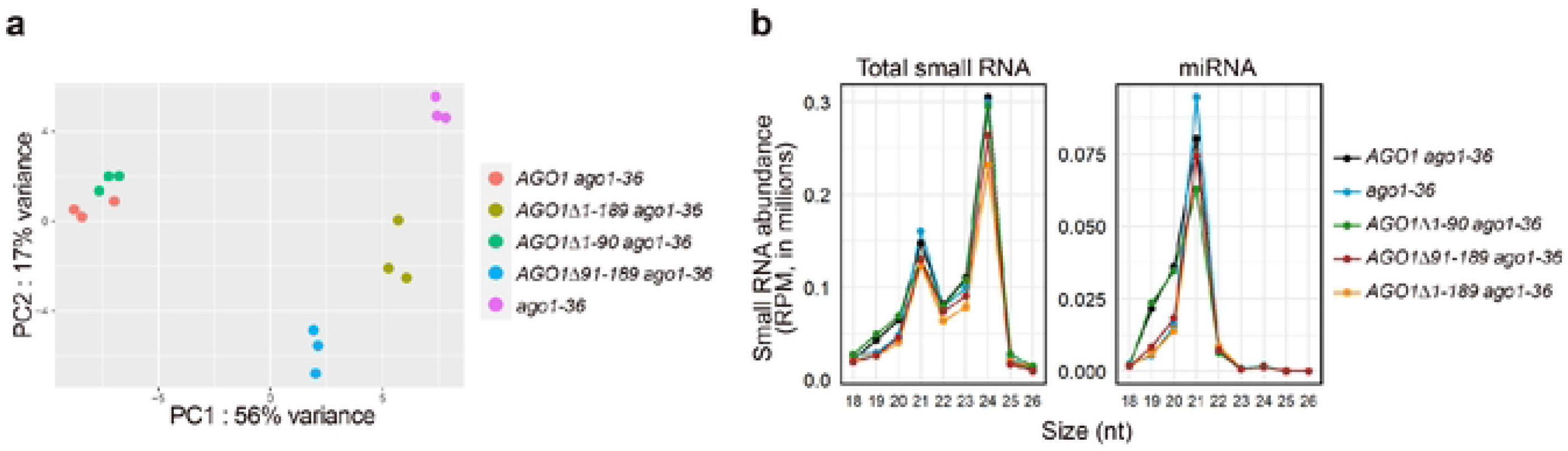
Analysis of small RNA seq data of AG01 NTE mutants. (a) PCA showing the reproducibility of the three replicates for each genotype. (b) Size (in nucleotides (nt)) distribution depicting the abundance of 18- to 26 nt total small RNAs and miRNAs in *ago1-36* expressing AG01 and AG01 NTE-truncated mutants. RPM, reads per million (see Methods).

**Figure S4.**
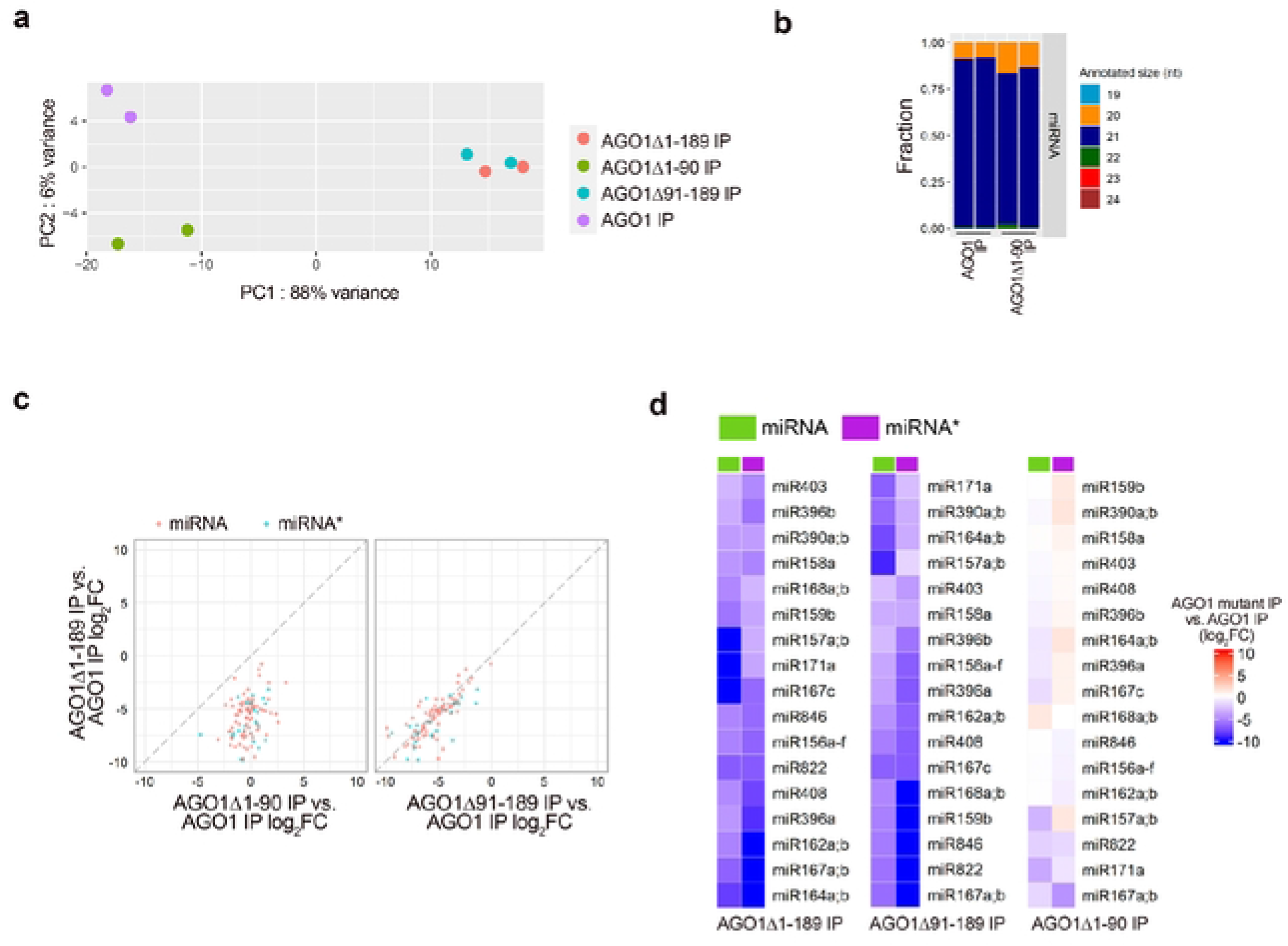
Analysis of small RNA-seq data of AG01 associated small RNAs (a) PCA analysis showing that the two biological replicates of each genotype were reproducible. (b) Bar plots showing the composit on of reads corresponding to annotated 19 to 24 nt miRNAs in AG01 IPs and AG01Δ1-90 IPs. Reads mapping to miRNA*s were excluded from this analysis. Annotated 20-nt miRNAs showed increased association with AG01Δ1-90 as compared to wild type AG01. (c) Scatter plots comparing the log_2_(fold change) of IP-ed miRNAs and miRNA*s between pairs of AG01 mutants. (d) Heatmap dep cting the levels of IP ed miRNAs and their corresponding miRNA*s in AG01 mutants. Note that only miRNAs for which the corresponding miRNA*s also passed the abundance filter (see Methods) are included in the heatmap.

**Figure S5.**
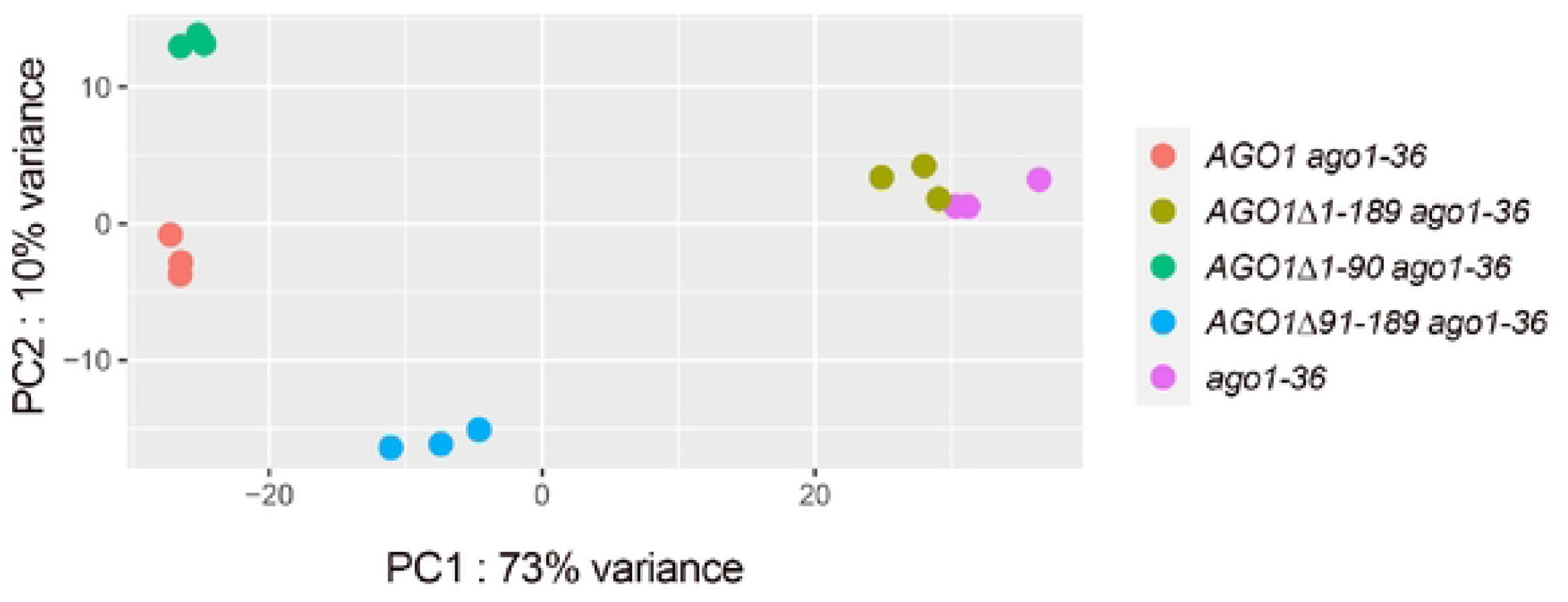
PCA analysis of RNA seq data of AG01 NTE mutants. Three biological replicates of each genotype cluster together

**Figure S6.**
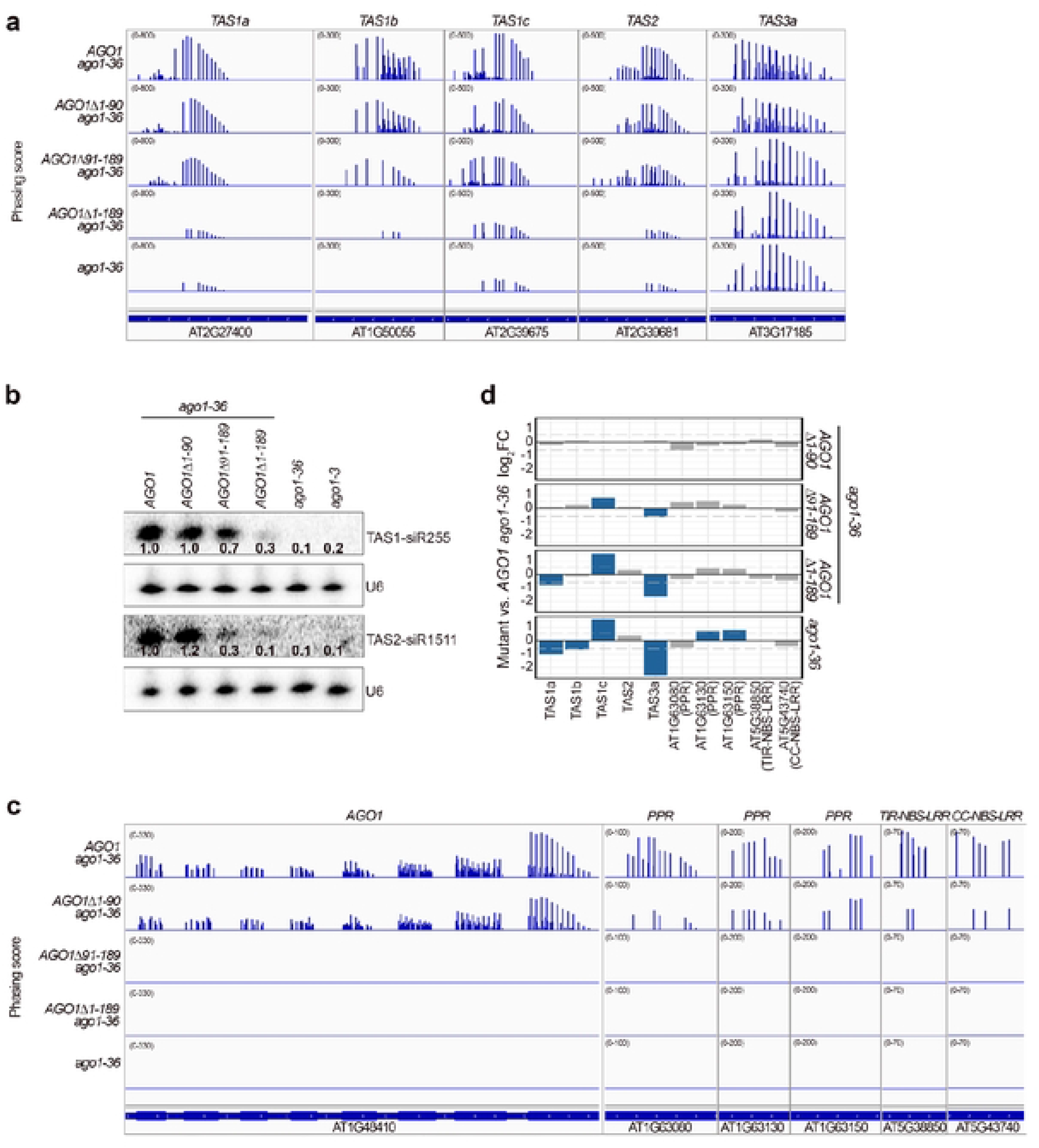
Biogenesis of phasiRNAs in NTE truncated AG01 mutants. (a) The phasing of small RNAs over the *TAS* transcripts in *ago1-36* and *ago 136* expressing wild-type *AG01* or *AG01* NTE mutants. (b) RNA gel blot analysis of ta siRNA abundance in 12 day old seedlings of *ago13, ago 136,* and *ago1-36* expressing *AG01* full-length or various NTE truncated forms. The numbers represent ta siRNA abundance in different genotypes relative to *AG01 agot-36.* The U6 blots serve as a loading control for the miRNA blots above. (c) The phasing of small RNAs over *AG01, PPR,* and *NBS-LRR* gene transcripts in *ago1-36* and *ago1-36* expressing wild-type *AG01* or *AG01* NTE mutants. (d) Bar plots showing the log_2_(fold change) of ta siRNAs or phasiRNAs from *TAS* and *PHAS* genes between *ago 136, ago1-36* expressing various *AG01* mutants and *AG01 ago1 36.* Blue bars denote ta siRNAs orphasiRNAs with sign ficantly d fferent expression (fold change > 1.5 and adjust *P* value < 0.01).

